# Development of rice conidiation media for *Ustilaginoidea virens*

**DOI:** 10.1101/641753

**Authors:** Yufu Wang, Fei Wang, Songlin Xie, Yi Liu, Jinsong Qu, Junbin Huang, Weixiao Yin, Chaoxi Luo

## Abstract

Rice false smut, caused by the ascomycete *Ustilaginoidea virens*, is a serious disease of rice worldwide. Conidia are very important infectious propagules of *U. virens*, but the ability of pathogenic isolates to produce conidia frequently decreases in culture, which influences pathogenicity testing. Here, we developed tissue media with rice leaves or panicles that stimulate conidiation of *U. virens*. Generally, rice leaf media more effectively increased conidiation than panicle media, and certain non-filtered tissue media were better than their filtered counterparts. Among the tested media, the Indica rice leaf medium with 0.06 g/ml of Wanxian 98 leaf was most efficient for inducing conidiation, and it was also usable for conidiation-defective isolates. Although the conidia induced in rice tissue media were smaller, they were able to germinate on potato sucrose agar medium and infect rice normally. This method provides a foundation for the production of conidia in *U. virens* that will be widely applied in the pathogenicity testing as well as in genetic analyses for false smut resistance in rice cultivars.

## Introduction

Rice false smut (RFS), caused by the ascomycete *Ustilaginoidea virens*, has become one of the most destructive diseases in the majority of rice-growing regions around the world, resulting in severe field losses of rice (Atia 2004; Ladhalakshmi et al. 2012; Rush et al. 2000; Sun et al. 2013; Zhou et al. 2008). The typical symptom is smut balls informed in rice panicles, the surface of which is covered by an abundance of powdery, dark-green chlamydospores (Ikegami 1962). Frequently, sclerotia can be observed on the surface of false smut balls. In addition to yield losses, *U. virens* produces ustiloxins that are toxic to human and animals and can inhibit radicle and plumule growth in plant seedlings (Abbas 2014; Meng et al. 2015; Wang et al. 2017).

*U. virens* can overwinter with thick-walled chlamydospores and sclerotia, the latter can produce sexual ascospores. Both ascospores and chlamydospores germinate and produce secondary conidia, which are the sources of primary infection in rice (Fan et al. 2014; Fan et al. 2010; Fu et al. 2012). Some studies have indicated that *U. virens* can infect roots and other rice tissues (Andargie 2015; Tang et al. 2017; Zheng et al. 2016). Nevertheless, numerous inoculation experiments and studies have shown that the majority of *U. virens* infections occur at the booting stage (Fu et al. 2012; Song et al. 2016; Tang et al. 2013; Yong et al. 2016). Conidia germinate and enter spikelets through the gap between the lemma and palea (Ashizawa et al. 2012), after that *U. virens* hyphae enter and grow intercellularly in the filaments of rice florets during the early stages of infection (Hu et al. 2014; Song et al. 2016; Tang et al. 2013). Subsequently, the hyphal growth extends to invade the stigmas and styles, enclosing the floral organs (Song et al. 2016). Finally, the hyphal mass extrudes from the lemma and palea, developing into false smut balls.

As one of the sources of primary infection, chlamydospores have been used to carry out artificial inoculation experiments; however, the incidence of infection achieved has been low (Fujita et al. 1989; Wang 1992; Zhang et al. 2003). In separate studies, mixtures of conidia and hyphae were injected into rice panicles at the booting stage, which can improve the incidence of infection significantly (Jia et al. 2015; Zhang et al. 2004; Zhang et al. 2003). Currently, this method is widely used to assess the pathogenicity of *U. virens* and the resistance of rice to RFS (Han et al. 2015; Li et al. 2013; Lv et al. 2016; Zheng et al. 2017; Zhou et al. 2014). The production of conidia depends on the time that the isolate has been in cultured and temperature, medium and isolates, etc. Up to 10^8^ conidia/ml can be produced after 6 days in potato dextrose broth (PDB) and potato sucrose broth (PSB) (Wang et al. 1998). Previous research has indicated that 2% sucrose-amended PDB containing barley seed produces more conidia in a shorter period of time (Ashizawa et al. 2011) compared with other media. The largest amount of conidia were detected after 6 or 7 days at 26-28°C with shaking at 140 to 210 rpm (He et al. 2011; Shi et al. 2017). In addition, some studies have found that conidia production was related to individual isolate characteristics (Li et al. 2012a; Li et al. 2012b). We also found that the conidiation capacity varies in different isolates and degradation with increasing numbers of transfers or, for certain isolates, length of time kept in the laboratory.

In this study, we developed tissue media with rice leaves or panicles that could promote the conidiation of *U. virens*. The optimal medium identified was especially useful for stimulating conidiation in isolates with defective conidial production. Although the conidia produced in this medium were small in size, they germinated and infected rice similarly with these produced in normal PSB medium.

## Materials and Methods

### *U. virens* isolates and rice cultivar

*U. virens* strains D32-1, HWD-2 and UV-8 as well as the GFP-labeled strain G2 were used in this study*. U. virens* isolates 09-11-1-1 and 09-14-21 were generously provided by Prof. Shiping Wang (Huazhong Agricultural University), both of which showed defective conidiation after being maintained in a laboratory environment for a long period of time (more than 8 years). The Indica rice cultivar Wanxian 98 and the Japonica rice cultivar Huajing 952 were selected for this study.

### Media and culture conditions

At the late booting stage (3-5 days before the heading stage of rice), leaves and panicles of the Indica and Japonica rice cultivars were collected for media preparation. Subsequently, 0.5, 1.5, 3.0 and 5.0 g of leaves or panicles of Wanxian 98 or Huajing 952 were crushed with a 22, 000 rpm mixer (Midea, model number: MJ-250BP01B) for 1 min in 40 ml distilled water respectively. Then they were used directly (unfiltered) or filtered with three layers of gauze to prepare media with the final concentrations of 0.01, 0.03, 0.06 and 0.10 g/ml respectively, and the media were autoclaved at 121°C for 30 min.

For conidiation tests, the strain G2 was incubated on PSA (potato sucrose agar: 200 ml of juice from 200 g of potato, 20 g of sucrose, and 15 g of agar per liter) at 27°C for 10 days, two 5-mm mycelial plugs were taken from the periphery of colony using a cork borer and incubated in 50 ml of PSB (PSA without agar) or liquid rice tissue media at 27°C with shaking at 160 rpm. The production of conidia was investigated from 3^rd^ to 7^th^ day using a hemacytometer. The conidiation from the filtered rice tissue media were compared with unfiltered ones at the 7^th^ day post-incubation (DPI). The conidiation in 0.06 g/ml leaf media were compared with 0.06 g/ml panicle media at 7 DPI. The conidiation in 0.06 g/ml of Indica rice leaves (IRLs) were compared with 0.06 g/ml of Japonica rice leaves (JRLs) at 7 DPI. Three replicates were observed for each treatment, with three droplets of each suspension examined per replicate. The experiment was conducted twice. The conidiation in different concentrations of rice tissue media and PSB were analyzed with SPSS at *P = 0.05*.

### Investigation of conidia size and germination and mycelial growth by *U. virens* strain

To determine conidia size, leaves or panicles from Wanxian 98 or Huajing 952 at late booting stage were collected to make 0.06 g/ml rice tissue media and PSB was selected as control. Strain G2 was shaken in liquid media for 7 days and filtered through four layers of gauze to collect its conidia. For each treatment, 100 conidia were measured at two perpendicular directions under a microscope. For the germination test, conidia were harvested by centrifugation at 7, 000 rpm and adjusted to a concentration of approximately 1.0×10^4^ conidia/ml. The conidia were spread on the surface of PSA or WA (water agar: 15 g of agar per liter), and the germination rate was investigated at 12, 24, 36 and 48 h respectively. For each treatment, 100 conidia were measured per replicate and three replicates were carried out. Each experiment was conducted twice. For mycelia growth, 5-mm mycelial plugs were removed from the margins of a colony and placed in the center of PSA, WA and 0.06 g/ml rice tissue media. The plates were incubated at 27°C in the dark for three weeks, and two perpendicular colony diameters were measured. Five replicates were performed for each treatment, and the experiment was conducted three times.

### Pathogenicity assays

After 7 days of incubation of strain G2 in PSB or 0.06 g/ml IRLs with shaking, conidia were collected and the concentration of each sample was adjusted to 1.0×10^6^ conidia/ml with PSB for inoculation or another treatment. For another treatment, the conidia were cultured in 50 ml of PSB at 27°C with shaking at 160 rpm for 8-12 h for germination, and then conidia were collected again and adjusted to 1.0×10^6^ conidia/ml with PSB for inoculation. The inoculation was performed as described previously (Song et al. 2016). In brief, approximately 2 ml of each conidia (1.0×10^6^ conidia/ml) suspension in PSB was injected into a single rice panicle at the late booting stage. Inoculated plants were kept in a 27°C greenhouse with 90-100% relative humidity (RH) for 7 days. Then, they were placed at 27°C and 80% RH until rice false smut symptoms appeared. The number of false smuts were recorded and analyzed. Twelve rice panicles were inoculated for each treatment, the experiments were conducted three times.

### Conidia production of conidiation-defective isolates in rice tissue media

The conidiation-defective isolates D32-1, HWD-2, UV-8, 09-11-1-1 and 09-14-21 were selected for conidiation tests, two 5-mm mycelial plugs were taken from the periphery of a 10-day-old colony of each isolate using a cork borer, and incubated in 50 ml of PSB or rice tissue media with the concentration of 0.06 g/ml IRL at 27°C with shaking at 160 rpm. The amounts of conidia were counted 3^rd^ to 7^th^ day using a hemacytometer. Three replicates were examined for each treatment, with three droplets of each suspension per replicate. The experiments were conducted twice.

## Results

### Rice tissue media promoted the conidiation of *U. virens*

To develop a medium that promotes *U. virens* conidiation, different rice tissue media were produced to investigate whether rice tissues can promote the conidiation of *U. virens*. Conidiation of strain G2 was evaluated at 3, 4, 5, 6 and 7 DPI. As shown in Table 1, compared with PSB, Indica rice leaves or panicles media produced more conidia at individual time points. These results indicated that Indica rice tissues could promote conidia production. Among the four concentrations, in general, higher rice tissue concentrations induced more conidia except for the medium with 0.10 g/ml IRLs, which showed fewer conidia compared with other concentrations of IRLs (Table 1). It is possible that too much fibrous tissue in the 0.10 g/ml IRL medium hindered the smooth shaking that is important for conidia production.

**Table 1.** Conidia production of *U. virens* strain G2 in Indica rice Wanxian 98 tissue media

| Medium<sup>y</sup> | Conidiation ($\times 10^6$ conidia/ml)<sup>x</sup> | | | | |
| --- | --- | --- | --- | --- | --- |
|  | 3 DPI | 4 DPI | 5 DPI | 6 DPI | 7 DPI |
| PSB | 0.000±0.000b | 0.022±0.022b | 0.367±0.367c | 1.544±0.953c | 1.911±0.946c |
| 0.01 g/ml IRL | 0.592±0.196a | 4.092±0.497a | 7.558±0.509b | 9.642±0.582b | 9.756±0.563b |
| 0.03 g/ml IRL | 0.253±0.185ab | 4.922±1.829a | 24.531±2.655a | 32.183±2.657a | 34.014±2.754a |
| 0.06 g/ml IRL | 0.094±0.039b | 1.092±0.505b | 11.144±3.621b | 32.931±3.847a | 38.964±4.741a |
| 0.10 g/ml IRL | 0.000±0.000b | 0.036±0.015b | 0.131±0.048c | 0.217±0.078c | 0.447±0.125c |
| PSB | 0.000±0.000b | 0.022±0.022b | 0.367±0.367b | 1.544±0.953d | 1.911±0.946d |
| 0.01 g/ml IRLF | 0.003±0.003b | 0.150±0.056a | 1.444±0.198b | 2.325±0.327d | 2.661±0.649d |
| 0.03 g/ml IRLF | 0.011±0.006b | 0.078±0.041a | 2.089±0.675b | 11.278±2.128c | 16.372±1.416c |
| 0.06 g/ml IRLF | 0.056±0.013a | 0.428±0.260a | 6.681±1.504a | 19.350±1.414b | 22.914±1.405b |
| 0.10 g/ml IRLF | 0.008±0.006b | 0.178±0.053a | 5.692±1.826a | 25.231±3.362a | 32.878±2.940a |
| PSB | 0.000±0.000b | 0.022±0.022b | 0.367±0.367b | 1.544±0.953c | 1.911±0.946d |
| 0.01 g/ml IRP | 1.303±0.462a | 2.689±0.682b | 3.089±0.520b | 3.639±0.541c | 3.675±0.633cd |
| 0.03 g/ml IRP | 0.283±0.159b | 2.336±0.677b | 5.667±1.202b | 8.389±1.858c | 7.939±1.056c |
| 0.06 g/ml IRP | 0.581±0.476ab | 7.939±2.135a | 27.236±3.534a | 27.683±2.080b | 28.853±1.305b |
| 0.10 g/ml IRP | 0.025±0.025b | 3.569±1.853b | 27.408±4.694a | 45.072±5.903a | 40.250±2.769a |
| PSB | 0.000±0.000b | 0.022±0.022b | 0.367±0.367c | 1.544±0.953c | 1.911±0.946c |
| 0.01 g/ml IRPF | 0.058±0.027a | 0.869±0.273ab | 1.903±0.383bc | 1.806±0.404c | 1.661±0.397c |
| 0.03 g/ml IRPF | 0.044±0.022ab | 0.839±0.499ab | 3.250±0.462b | 4.633±1.171c | 4.536±0.620c |
| 0.06 g/ml IRPF | 0.044±0.025ab | 2.925±1.925a | 10.397±0.544a | 24.253±1.071a | 27.544±1.113a |
| 0.10 g/ml IRPF | 0.006±0.006ab | 0.328±0.153ab | 7.917±1.917a | 17.228±2.699b | 21.288±2.107b |

<sup>x</sup> Data shown are the means of two independent experiments, reported as the mean±standard error. Values followed by the same letters within the same column for the same media with different concentrations of rice tissue are not significantly different based on one-way ANOVA with LSD tests performed with SPSS at $P = 0.05$ . Data were logarithm-transformed before analysis. The same PSB culture was used as the control for the different rice tissue media.
DPI: days post incubation.
<sup>y</sup> PSB: potato sucrose broth; IRL: Indica rice leaf; IRLF: Indica rice leaf filtrate; IRP: Indica rice panicle; IRPF: Indica rice panicle filtrate.

The effect of different media with unfiltered or filtered rice leaves and panicles was evaluated. The number of conidia from IRL media was compared to that from IRL filtrate (IRLF) media at 7 DPI. Results showed that more conidia were produced in IRL media at concentrations of 0.01 (*P* = 8.873×10^−6^), 0.03 (*P* = 7.383×10^−4^) and 0.06 g/ml (*P* = 1.755×10^−2^). However, at 0.10 g/ml, fewer conidia were produced in the IRL medium compared with the IRLF medium, most likely because too much rice leaf tissue was present (*P* = 1.071×10^−4^). The similar results were also observed at 3, 4, 5 and 6 DPI.

Considering that *U. virens* infects the rice panicle, panicle tissues were also selected to prepare media for conidiation testing. The results of the Indica rice panicle (IRP) media were similar to these of the IRL media at 7 DPI, with the exception of the 0.10 g/ml IRP medium that induced more conidia production (*P* = 4.061×10^−4^), likely because there is less fibrous tissue in rice panicles compared to leaves. Four concentrations of IRPs and IRP filtrates (IRPFs) were also compared. More conidia were observed in the IRPs media, except for 0.06 g/ml media which did not show significant difference between the IRP and IRPF media (*P* = 0.463).

For the different tissues, compared with IRP, the IRL media produced more conidia at 0.01 (*P* = 3.218×10^−5^) and 0.03 g/ml (*P* = 7.818×10^−5^) at 7 DPI, but no significant difference was observed at 0.06 g/ml (*P* = 0.086). However, at 0.10 g/ml, the IRLs media produced fewer conidia compared with the IRP media (*P* = 2.867×10^−5^).

Similar results were observed when tissues of Japonica rice cultivar Huajing 952 were used. Generally, the rice tissue media produced more conidia, except for the 0.01 g/ml Japonica rice leaf filtrate (JRLF) medium. For the different tissues, at 0.06 g/ml, Japonica rice leaf (JRL) medium produced more conidia than Japonica rice panicle (JRP) medium at 7 DPI (*P* = 2.459×10^−2^); however, no significant differences were observed at other concentrations. Compared with JRLF media, JRL media produced more conidia at 0.01 (*P* = 4.608×10^−4^), 0.03 (*P* = 2.464×10^−5^) and 0.06 g/ml (*P* = 6.709×10^−7^), and no significant difference was observed at 0.10 g/ml (*P* = 0.340). JRP media produced more conidia at all four concentrations than JRPF media (Table 2).

**Table 2.** Conidia production of *U. virens* strain G2 in Japonica rice Huajing 952 tissue media

| Medium<sup>y</sup> | Conidiation ($\times 10^6$ conidia/ml)<sup>x</sup> | | | | |
| --- | --- | --- | --- | --- | --- |
|  | 3 DPI | 4 DPI | 5 DPI | 6 DPI | 7 DPI |
| PSB | 0.000±0.000b | 0.022±0.022c | 0.367±0.367d | 1.544±0.953d | 1.911±0.946e |
| 0.01 g/ml JRL | 0.486±0.128a | 3.347±0.359a | 5.544±0.196c | 6.886±0.580c | 7.275±0.774d |
| 0.03 g/ml JRL | 0.097±0.032b | 2.442±0.578ab | 14.847±1.002b | 25.942±1.584b | 25.252±1.326b |
| 0.06 g/ml JRL | 0.103±0.033b | 2.236±0.461b | 31.794±2.394a | 38.722±2.351a | 36.789±1.511a |
| 0.10 g/ml JRL | 0.053±0.028b | 0.467±0.183c | 2.128±0.744cd | 5.050±1.199cd | 12.150±0.898c |
| PSB | 0.000±0.000b | 0.022±0.022b | 0.367±0.367b | 1.544±0.953b | 1.911±0.946b |
| 0.01 g/ml JRLF | 0.011±0.008b | 0.206±0.134b | 0.758±0.196b | 1.658±0.069b | 1.778±0.193b |
| 0.03 g/ml JRLF | 0.081±0.023a | 2.608±0.690a | 4.769±0.663a | 5.850±0.562a | 7.989±1.739a |
| 0.06 g/ml JRLF | 0.086±0.024a | 0.583±0.229b | 4.361±1.184a | 8.961±2.533a | 10.550±1.851a |
| 0.10 g/ml JRLF | 0.058±0.024ab | 0.392±0.174b | 5.022±2.004a | 9.603±1.581a | 10.561±1.296a |
| PSB | 0.000±0.000b | 0.022±0.022c | 0.367±0.367d | 1.544±0.953d | 1.911±0.946d |
| 0.01 g/ml JRP | 0.333±0.187a | 1.161±0.681bc | 2.456±0.824d | 3.219±1.004d | 4.022±1.061d |
| 0.03 g/ml JRP | 0.075±0.024ab | 5.203±0.763a | 12.036±0.685c | 11.697±0.715c | 13.156±1.364c |
| 0.06 g/ml JRP | 0.275±0.155ab | 2.767±0.789b | 19.103±4.037b | 18.719±2.413b | 26.008±3.464b |
| 0.10 g/ml JRP | 0.081±0.047ab | 2.983±0.621b | 34.817±3.336a | 49.564±3.602a | 52.244±2.951a |
| PSB | 0.000±0.000b | 0.022±0.022b | 0.367±0.367c | 1.544±0.953c | 1.911±0.946c |
| 0.01 g/ml JRPF | 0.025±0.012a | 0.228±0.104b | 0.531±0.109c | 0.619±0.128c | 0.864±0.180c |
| 0.03 g/ml JRPF | 0.014±0.009ab | 0.236±0.073b | 1.364±0.321bc | 2.789±0.328c | 4.308±0.384c |
| 0.06 g/ml JRPF | 0.008±0.008ab | 0.358±0.077b | 6.653±0.811b | 11.139±1.868b | 14.092±0.484b |
| 0.10g/ml JRPF | 0.022±0.006ab | 1.686±0.407a | 12.628±3.970a | 27.444±4.205a | 26.050±3.017a |

<sup>x</sup> Data shown are the means of two independent experiments, reported as the mean±standard error. Values followed by the same letters within the same column for the same media with different concentrations of rice tissue are not significantly different based on one-way ANOVA with least significant difference performed with SPSS at $P = 0.05$ . Data were logarithm-transformed before analysis. The PSB was used as the control for the different rice tissue media.
DPI: days post incubation.
<sup>y</sup> PSB: potato sucrose broth; JRL: Japonica rice leaf; JRLF: Japonica rice leaf filtrate; JRP: Japonica rice panicle; JRPF: Japonica rice panicle filtrate.

The conidia from 0.06 g/ml rice leaf media at 7 DPI were compared between rice cultivars Wanxian 98 and Huajing 952, and no significant difference was observed between IRL and JRL media (*P* = 0.677). To compare among different growth stage leaves, the first and second leaves from the top of rice cultivar Wanxian 98 at three or four days prior to the heading stage as well as leaves at the tillering stage were collected and used to make 0.06 g/ml media. No significant difference was observed among the different types of leaves (*P* = 0.795).

### Smaller conidia were produced in the rice tissue media

To investigate whether the conidia produced in the rice tissue media were similar to these produced in PSB, the conidial size (length and width) of strain G2 were investigated. Conidia produced in PSB and IRL, IRP, JRL and JRP media were collected, and the length and width were measured. The results showed that the conidia produced in PSB were larger than these produced in IRL, IRP, JRL and JRP media, with these produced in the IRP media being the smallest (Table 3). These results indicated that even though the rice tissue media could stimulate the conidiation of *U. virens*, the conidia produced were smaller than these produced in PSB.

**Table 3.** Size of conidia of *U. virens* strain G2 produced in PSB and rice tissue media^x^

|  | PSB<sup>y</sup> | IRL | IRP | JRL | JRP |
| --- | --- | --- | --- | --- | --- |
| Length (μm) | 5.895±0.109a | 5.017±0.087b | 4.706±0.064c | 5.166±0.078b | 5.114±0.056b |
| Width (μm) | 3.395±0.047a | 3.262±0.032b | 3.195±0.032b | 3.090±0.028c | 3.007±0.026c |

<sup>x</sup> Data shown are the means of the size of 100 conidia, reported as the mean±standard deviation. Values followed by the same letters within the same row are not significantly different based on one-way ANOVA with least significant difference tests performed with SPSS at $P = 0.05$ .
<sup>y</sup> PSB: potato sucrose broth; IRL: Indica rice leaf medium; IRP: Indica rice panicle medium; JRL: Japonica rice leaf medium; JRP: Japonica rice panicle medium.

### Germination was delayed for the conidia produced in rice tissue media

To investigate the germination of conidia produced in rice tissue media, conidial germination of strain G2 from IRL, IRP, JRL and JRP media as well as PSB were assessed on PSA and WA at 12, 24, 36 and 48 h. On PSA, the germination rate of conidia from PSB was 96.9% at 12 h. For conidia from IRL and JRL media, the germination rates were 68.9% and 67.8%, respectively. Conidia from IRP and JRP media showed the highest (89.9%) and lowest (41.6%) germination rates among these from the four different rice tissue media (Fig 1). After more than 24 h, almost all conidia had germinated. On WA, a germination rate of 43.6% was observed for conidia from PSB at 12 h, while lower germination rates were found for conidia from other media. Over time, the germination rate increased gradually and more than 90% of conidia from PSB germinated at 48 h. However, the germination rates were 69.2%, 56.1%, 54.2% and 37.7% for conidia from IRL, IRP, JRL and JRP media, respectively, which were significantly lower than that from PSB (Fig 1). These results showed that the germination of conidia from rice tissue medium was slower than that from PSB, and generally the germination rate was higher on PSA than on WA.

**Fig 1.**
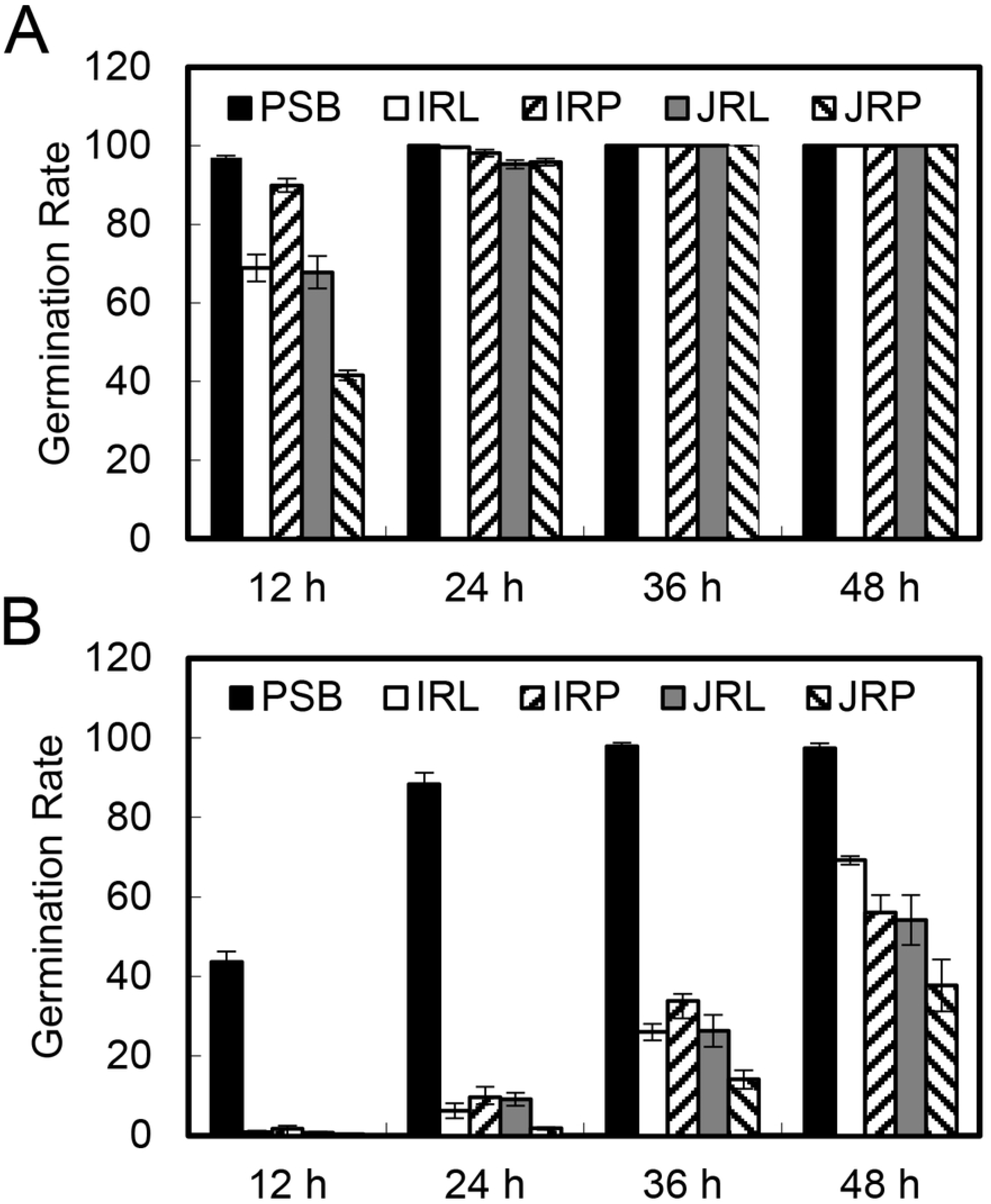
Conidia produced in rice tissue media have decreased germination rates. Conidia produced in PSB, IRL, IRP, JRL and JRP media were spread on PSA (**A**) or WA (**B**) and their germination rates were measured at 12 h, 24 h, 36 h and 48 h. Three independent experiments were performed, and similar results were obtained. Error bars represent the standard deviation of three replicates.

### The mycelial mass was thin on rice tissue media

Considering that rice tissue media promotes the production of conidia, we investigated whether the media provide enough nutrients for *U. virens* growth. Mycelial growth of strain G2 was assessed on rice tissue media solidified with 2% agar. The results showed that the colony diameter was largest on JRP media with agar (JRPA), while they were smaller on PSA and other rice tissue media. On WA, though some hyphae were observed, they grew slowly and loosely. Basically, despite the fact that mycelial growth was stimulated on some rice tissue media, the hyphae were loose and mycelia were thin (Fig 2). These results showed that *U. virens* can grow on both rice tissue media and PSA, but the overall biomass is decreased on rice tissue media, suggesting that rice tissue media could not provide enough nutrients for mycelial growth.

**Fig 2.**
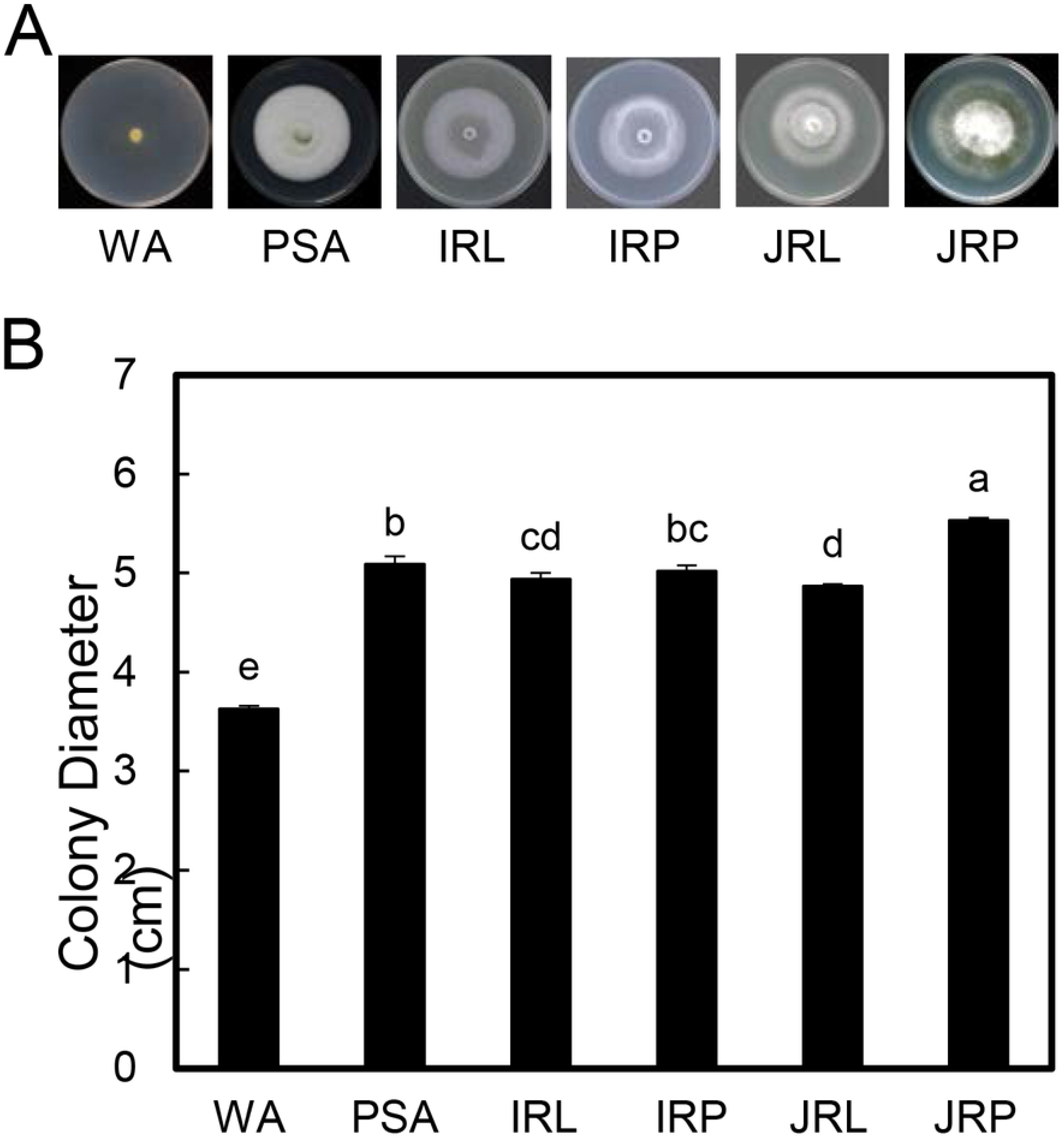
Rice tissue media do not affect hyphal growth. Plugs of strain G2 were inoculated onto WA, PSA, JRL and JRP media. The colonies were photographed (**A**), and their diameters (**B**) were measured for 3 weeks after inoculation. Three independent experiments were performed, and similar results were obtained. Error bars represent the standard deviation of three replicates, and the different letters above each column indicate statistical significance (*P* < 0.05).

### Pathogenicity test for the induced condinia

To investigate the pathogenicity of conidia produced in rice tissue media, conidia of strain G2 from 0.06 g/ml IRL medium were collected for inoculation. Three treatments were performed: (I) the conidia were collected and diluted with water, (II) the conidia were collected and diluted with PSB, and (III) the conidia were collected and cultured in PSB for 8-12 h, then collected again and diluted with PSB. The inoculation results showed that more smut balls were observed for treatment III, whereas few or no smut balls were observed for treatments I and II (Table 4). These results showed that although the conidia produced in rice tissue media can germinate and infect rice successfully, they should be incubated for 8-12 h in PSB before they are used to inoculate rice.

**Table 4.** Pathogenicity testing of the different conidia treatments

| Treatment<sup>x</sup> | NO. of false smuts<sup>y</sup> |
| --- | --- |
| I | 0.5±0.07b |
| II | 0.1±0.3b |
| III | 20.9±10.5a |

<sup>x</sup> Conidia from *U. virens* strain G2 from PSB and 0.06 g/ml IRLs media at 7 DPI were collected, and then treated as follow: I, the conidia were collected and diluted with water; II, the conidia were collected and diluted with PSB; III, the conidia were collected and cultured in PSB for 8-12 h and collected again and diluted with PSB.
<sup>y</sup> Data shown are the means of three independent experiments, reported as the mean±standard error. Values not followed by the same letter within a column differ significantly based on one-way ANOVA with least significant difference tests performed with SPSS at $P = 0.05$ .

### IRL induced conidiation in conidiation-defective isolates

As the best medium to induce conidiation among examined different media, the 0.06 g/ml IRL medium was used to investigate whether conidiation could be induced in conidiation-defective isolates. The isolates D32-1, HWD2-2, UV-8, 09-11-1-1 and 09-14-21 were selected because they all fail to produce conidia or only produce few conidia. Interestingly, for all of these isolates, conidia were produced at 3 or 4 DPI in 0.06 g/ml IRL medium, with substantial amounts of conidia produced at 6 or 7 DPI; conversely, little or no conidia were observed in PSB. HWD2-2 and UV-8 produced the most conidia (up to 2.3×10^7^ conidia/ml) in the IRL medium, an amount equal to that produced by non-defective isolates in PSB. The other isolates, D32-1, 09-11-1-1 and 09-14-21, produced fewer conidia but were still within the same order of magnitude as that produced by HWD2-2 and UV-8. These results indicated that the IRL medium could be used to induce conidiation in defective isolates (Table 5).

**Table 5.**
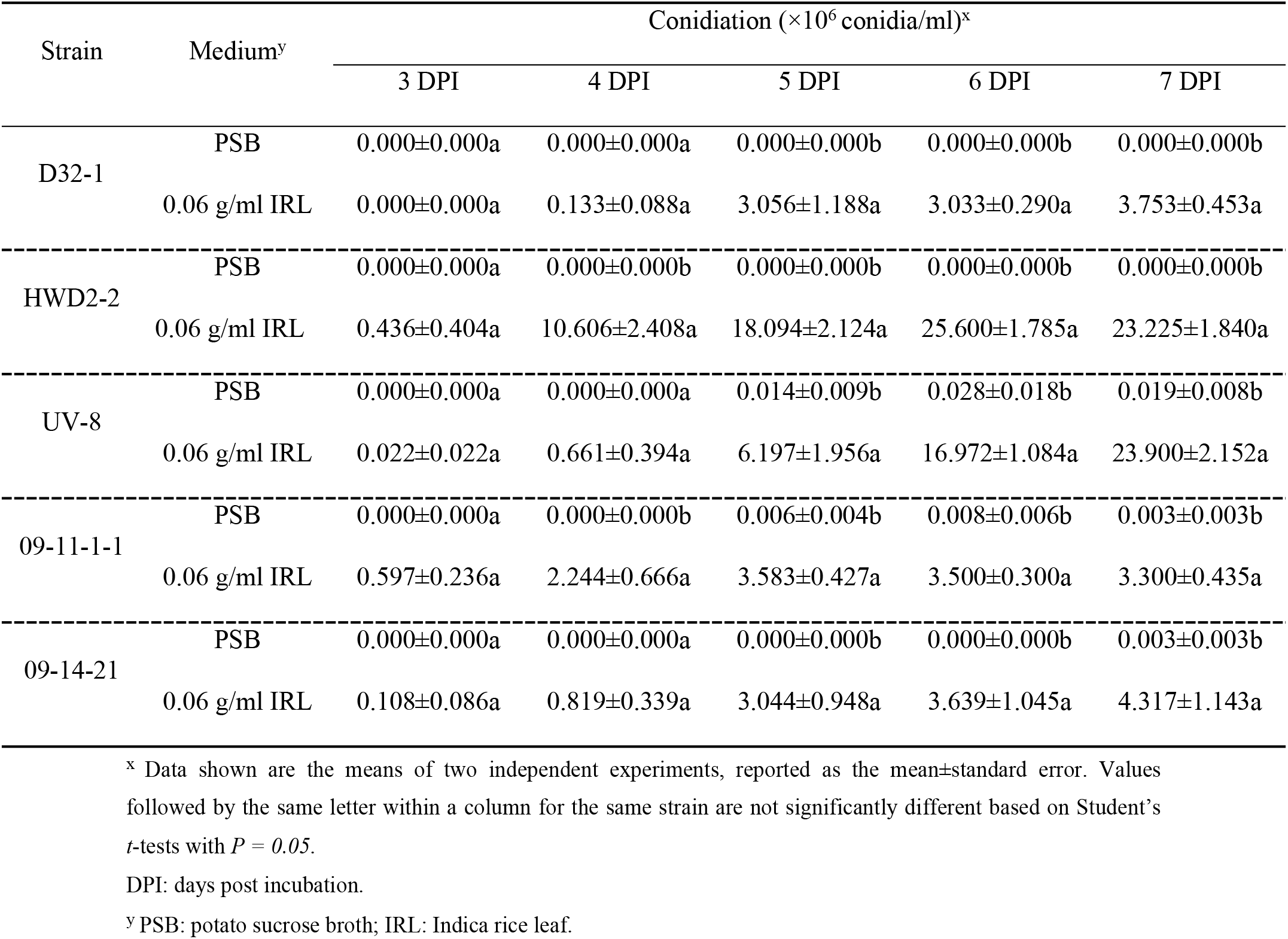
Conidiation of defective isolates in Wanxian 98 rice leaf medium

## Discussion

In this study, we developed rice tissue media that could promote the conidiation of *U. virens*. These media were especially useful for the production of conidia in conidiation-defective strains. Even though the conidia produced in rice tissue media were smaller and germinated more slowly compared with these produced in PSB, the pathogenicity of the tested strain has not been effected once they were incubated in PSB for 8-12 h before inoculation, indicating that rice tissue media, especially the 0.06 g/ml IRL can be used to provide conidia for pathogenicity testing or genetic analyses of rice resistance and *U. virens* avirulence.

The major factors that affect conidiation include nutrients, host tissues and light (Su et al. 2012). For *U. virens*, temperature, medium and strain are important for conidia production. PSB and PDB media were commonly used for conidia production (Li et al. 2012a; Shi et al. 2017; Wang et al. 1998). However, conidiation varies among different isolates and conidia production can be unstable, eventually receding along with the increased numbers of transfer or long-term storage. Host tissues are often used to induce sporulation in plant pathogenic fungi (Su et al. 2012). Research has shown that different kinds of plant leaves can induce sporulation in *Colletotrichum dematium* (Yoshida and Shirata 2000). Furthermore, the biotin in mulberry leaves was found to alter the synthesis of cell wall polysaccharides and oleic acid, thus triggering the selective expression of genes involved in sporulation (Yoshida and Shirata 2000). A carrot medium was considered as the best medium for obtaining conidia from *Venturia nashicola* (Choi et al. 2017). In addition, dried barley seeds were added to 2% sucrose-amended PDB medium to improve conidia production in *U. virens* (Ashizawa et al. 2011). Starvation or nutritional depletion often stimulates sporulation (Skromne et al. 1995), but no clear research has been performed for *U. virens*. In this study, rice leaves and panicles were crushed to make media in which they induced conidiation within a shorter timeframe than the basal medium. No significant differences were observed between the leaves and panicles from Indica cultivar Wanxian 98. However, for Japonica cultivar Huajing 952, media containing leaves produced more conidia than panicles. No differences were observed between leaf media and leaves from different varieties or stages of maturity. The conidiation capacity of *U. virens* on different media was positively related to tissue concentrations, except for 0.1 g/ml IRL in which insufficient shaking occurred. The media with rice leaf or panicle filtrates produced fewer conidia compared with unfiltered media, probably because the residues contain substances or provide certain physical benefits that affect conidiation. Meanwhile, less conidia were produced in 0.01 g/ml JRPF than in PSB, which might have been caused by insufficient nutrients. All these rice tissue contain less nutrition compared with PSB, indicating appropriate starvation could stimulate the conidiation. On the other side, some substance in rice may play the function to promote the conidiation.

Conidia can germinate on biotic and abiotic surfaces to form hyphae and differentiate conidiophores to generate a large number of secondary conidia or produce secondary conidia directly (Fan et al. 2014; Fu et al. 2012). The conidia produced in rice tissue media were smaller with slower germination than these produced in PSB, suggesting that rice tissue media does not supply enough nutrients for the conidial growth. On WA, the germination rates of the conidia from rice tissue media were lower than these produced in PSB, indicating that sufficient nutrition is important for germination of conidia.

Conidia produced in rice leaf media and then cultured in PSB for 8-12 h with shaking were able to infect rice. However, conidia produced without cultured in PSB lost the ability to infect rice, even they were diluted in PSB. For successful inoculation, the conidia have to be diluted or resuspended in PSB (Jia et al. 2015; Zhang et al. 2004). These results indicated that conidia produced in rice tissue media do not have enough nutrition and should be further cultured in PSB to achieve successful infection.

In conclusion, we developed media that induced conidiation in *U. virens*, even in conidiation-defective isolates. These media are extremely valuable because many important isolates frequently lose their ability to conidiate after many transfers or long-term storage. Our media could stimulate and recover conidiation in such isolates, thus providing a solid foundation for continuously performing pathogenicity testing, which is necessary for genetic analyses of rice resistance and fungal virulence test.

## Author Contributions

Conceived and designed the experiments: YW, FW, WY and CL. Performed the experiments: YW, SX, FW, YL and JQ. The data analysis: YW, WY, JH and CL. The article Writing: YW, WY, JH and CL.

## References

Abbas, H., Shier, W., Cartwright, R. and Sciumbato, G. 2014. *Ustilaginoidea virens* infection of rice in arkansas toxicity of false smut galls, their extracts and the ustiloxin fraction. American Journal of Plant Sciences 5:3166–3176.

Andargie, M., Li, L., Feng, A. and Li, J. 2015. Colonization of rice roots by a green fluorescent protein-tagged isolate of *Ustilaginoidea virens*. American Journal of Plant Sciences 6:2272–2279.

Ashizawa, T., Takahashi, M., Moriwaki, J., and Hirayae, K. 2011. A refined inoculation method to evaluate false smut resistance in rice. J Gen Plant Pathol 77:10–16.

Ashizawa, T., Takahashi, M., Arai, M., and Arie, T. 2012. Rice false smut pathogen, *Ustilaginoidea virens*, invades through small gap at the apex of a rice spikelet before heading. J Gen Plant Pathol 78:255–259.

Atia, M. M. M. 2004. Rice false smut (*Ustilaginoidea virens*) in Egypt. Journal of Plant Diseases and Protection 111:71–82.

Choi, E. D., Kim, G. H., Lee, Y. S., Jung, J. S., Song, J. H., and Koh, Y. J. 2017. Development of carrot medium suitable for conidia production of *Venturia nashicola*. Plant Pathology J 33:75–79.

Fan, J., Guo, X. Y., Huang, F., Li, Y., Liu, Y. F., Li, L., Xu, Y. J., Zhao, J. Q., Xiong, H., Yu, J. J., and Wang, W. 2014. Epiphytic colonization of *Ustilaginoidea virens* on biotic and abiotic surfaces implies the widespread presence of primary inoculum for rice false smut disease. Plant Pathology 63:937–945.

Fan, R., Wang, Y., Liu, B., Zhang, J., Wang, H., and Hu, D. 2010. The process of asexual spore formation and examination of chlamydospore germination of *Ustilaginoidea virens*. Mycosystema 29:188–192. In Chinese, abstract in English.

Fu, R., Ding, L., Zhu, J., Li, P., and Zheng, A. P. 2012. Morphological structure of propagules and electrophoretic karyotype analysis of false smut *Villosiclava virens* in rice. Journal of microbiology 50:263–269.

Fujita, Y., Sonoda, R., and Yaegashi, H. 1989. Inoculation with conidiospores of false smut fungus to rice panicles at the booting stage. Ann. Phytopath. Soc. Japan 55:629–634.

Han, Y., Zhang, K., Yang, J., Zhang, N., Fang, A., Zhang, Y., Liu, Y., Chen, Z., Hsiang, T., and Sun, W. 2015. Differential expression profiling of the early response to *Ustilaginoidea virens* between false smut resistant and susceptible rice varieties. BMC Genomics 16:955.

He, H., Chen, X., Yang, X., Wu, S., Wang, L., Wongkaew, S., and Yuan, J. 2011. Isolation technology of single spore and optimization of conidium culture condition in rice *Ustilaginoidea virens*. Guizhou Agricultural Sciences 39:119–121. In Chinese, abstract in English.

Hu, M. L., Luo, L. X., Wang, S., Liu, Y. F., and Li, J. Q. 2014. Infection processes of *Ustilaginoidea virens* during artificial inoculation of rice panicles. Eur J Plant Pathol 139:67–77.

Ikegami, H. 1962. Studies on the false smut of rice. V. Seedling inoculation with the chlamydospores of the false smut fungus. Japanese Journal of Phytopathology 27:16–23.

Jia, Q., Lv, B., Guo, M. Y., Luo, C. X., Zheng, L., Hsiang, T., and Huang, J. B. 2015. Effect of rice growth stage, temperature, relative humidity and wetness duration on infection of rice panicles by *Villosiclava virens*. Eur J Plant Pathol 141:15–25.

Ladhalakshmi, D., Laha, G. S., Singh, R., Karthikeyan, A., Mangrauthia, S. K., Sundaram, R. M., Thukkaiyannan, P., and Viraktamath, B. C. 2012. Isolation and characterization of *Ustilaginoidea virens* and survey of false smut disease of rice in India. Phytoparasitica 40:171–176.

Li, W., Li, L., Feng, A., Zhu, X., and Li, J. 2013. Rice false smut fungus, *Ustilaginoidea virens*, inhibits pollen germination and degrades the integuments of rice ovule. American Journal of Plant Sciences 4:2295–2304.

Li, Y., Yin, X., Liu, Y., Yu, J., and Chen, Z. 2012a. Relativity of biological characteristics and pathogenicity of *Ustilaginoidea virens*. Acta Phytopathologica Sinica 42:353–364. In Chinese, abstract in English.

Li, Y., Yu, J., Liu, Y., Yin, X., Zhang, R., Yu, M., and Chen, Z. 2012b. Determination of sporulation and pathogenicity of *Ustilaginoidea virens*. Scientia Agricultura Sinica 45:4166–4177. In Chinese, abstract in English.

Lv, B., Zheng, L., Liu, H., Tang, J. T., Hsiang, T., and Huang, J. B. 2016. Use of random T-DNA mutagenesis in identification of gene *UvPRO1*, a regulator of conidiation, stress response, and virulence in *Ustilaginoidea virens*. Front. Microbiol 7:2086.

Meng, J., Sun, W., Mao, Z., Xu, D., Wang, X., Lu, S., Lai, D., Liu, Y., Zhou, L., and Zhang, G. 2015. Main ustilaginoidins and their distribution in rice false smut balls. Toxins 7:4023–4034.

Rush, M. C., Shahjahan, A. K. M., Jones, J. P., and Groth, D. E. 2000. Outbreak of false smut of rice in Louisiana. Plant Dis 84:100–100.

Shi, T., Wang, Z., Cai, C., Yang, P., Qin, Q., and Zhang, J. 2017. Study on factors influencing prepration of thin-walled conidia of *Ustilaginoidea virens*. Plant Protection 43:131–134. In Chinese, abstract in English.

Skromne, I., Sanchez, O., and Aguirre, J. 1995. Starvation stress modulates the expression of the *Aspergillus nidulans brlA* regulatory gene. Microbiology 141 (Pt 1):21–28.

Song, J. H., Wei, W., Lv, B., Lin, Y., Yin, W. X., Peng, Y. L., Schnabel, G., Huang, J. B., Jiang, D. H., and Luo, C. X. 2016. Rice false smut fungus hijacks the rice nutrients supply by blocking and mimicking the fertilization of rice ovary. Environmental microbiology 18:3840–3849.

Su, Y.-Y., Qi, Y.-L., and Cai, L. 2012. Induction of sporulation in plant pathogenic fungi. Mycology 3:195–200.

Sun, X., Kang, S., Zhang, Y., Tan, X., Yu, Y., He, H., Zhang, X., Liu, Y., Wang, S., Sun, W., Cai, L., and Li, S. 2013. Genetic diversity and population structure of rice pathogen *Ustilaginoidea virens* in China. PLoS One 8:e76879.

Tang, J. T., Zheng, L., Jia, Q., Liu, H., Hsiang, T., and Huang, J. B. 2017. PCR markers derived from comparative genomics for detection and identification of the rice pathogen *Ustilaginoidea virens* in plant tissues. Plant Dis 101:1515–1521.

Tang, Y. X., Jin, J., Hu, D. W., Yong, M. L., Xu, Y., and He, L. P. 2013. Elucidation of the infection process of *Ustilaginoidea virens* (teleomorph: *Villosiclava virens*) in rice spikelets. Plant Pathology 62:1–8.

Wang, G. 1992. Studies on the infection period and the infection gate of the chlamydospores of *Ustilaginoidea virens* (Cooke)TAK. on rice. Acta Phytophylacica Sinica 19:97–100. In Chinese, abstract in English.

Wang, S., Bai, Y., Zhou, Y., Yao, J., and Bai, J. 1998. The pathogen of false smut of rice. Acta Phytopathologica Sinica 28:19–24. In Chinese, abstract in English.

Wang, X., Wang, J., Lai, D., Wang, W., Dai, J., Zhou, L., and Liu, Y. 2017. Ustiloxin G, a new cyclopeptide mycotoxin from rice false smut balls. Toxins 9:54.

Yong, M. L., Fan, L. L., Li, D. Y., Liu, Y. J., Cheng, F. M., Xu, Y., Wang, Z. Y., and Hu, D. W. 2016. *Villosiclava virens* infects specifically rice and barley stamen filaments due to the unique host cell walls. Microscopy research and technique 79:838–844.

Yoshida, S., and Shirata, A. 2000. Biotin induces sporulation of mulberry anthracnose fungus, *Colletotrichum dematium*. J. Gen. Plant Pathol 66:117–122.

Zhang, J., Zhang, B., Chen, Z., Liu, Y., and Lu, F. 2003. Preliminary study on inoculation method of rice false smut and its effect. Chinese J Rice Sci 17:390–392. In Chinese, abstract in English.

Zhang, J., Chen, Z., Zhang, B., Liu, Y., and Lu, F. 2004. Inoculation techniques used for inducing Rice false smut efficiently. Acta Phytopathologica Sinica 34:463–467. In Chinese, abstract in English.

Zheng, D. W., Zhang, Y. M., Yin, T., Xu, J. R., and Wang, C. F. 2016. Molecular detection of *Ustilaginoidea virens* in rice plant. Acta Phytopathologica Sinica 46:145–150. In Chinese, abstract in English.

Zheng, M. T., Ding, H., Huang, L., Wang, Y. H., Yu, M. N., Zheng, R., Yu, J. J., and Liu, Y. F. 2017. Low-affinity iron transport protein Uvt3277 is important for pathogenesis in the rice false smut fungus *Ustilaginoidea virens*. Curr Genet 63:131–144.

Zhou, Y. L., Pan, Y. J., Xie, X. W., Zhu, L. H., Xu, J. L., Wang, S., and Li, Z. K. 2008. Genetic diversity of rice false smut fungus, *Ustilaginoidea virens* and its pronounced differentiation of populations in North China. J Phytopathology 156:559–564.

Zhou, Y. L., Xie, X. W., Zhang, F., Wang, S., Liu, X. Z., Zhu, L. H., Xu, J. L., Gao, Y. M., and Li, Z. K. 2014. Detection of quantitative resistance loci associated with resistance to rice false smut (*Ustilaginoidea virens*) using introgression lines. Plant Pathology 63:365–372.

